# A CRISPR/Cas9 mediated 53 kb deletion of the NRC4 gene cluster of tomato does not affect bacterial flagellin-triggered immunity

**DOI:** 10.1101/697425

**Authors:** Chih-Hang Wu, Hiroaki Adachi, Juan Carlos De la Concepcion, Roger Castells-Graells, Vladimir Nekrasov, Sophien Kamoun

## Abstract

Plants utilise cell surface pattern recognition receptors (PRRs) and intracellular nucleotide-binding domain leucine-rich repeat containing receptors (NLRs) to fend off invading pathogens. Although PRR- and NLR-triggered immunity are generally thought to activate distinct pathways, they can induce similar outputs. However, whether these two pathways converge at some point to potentiate and strengthen the immune response remains unclear. For instance, the extent to which the tomato NLR helper NRC4 is implicated in response to the bacterial flagellin peptide flg22 needs to be elucidated. One challenge is that the tomato NRC4 gene cluster consists of three paralogues and the related NRC5 gene. Here, we took advantage of the CRISPR/Cas9 system to generate a tomato mutant with a 53 kb deletion that encompasses the four NRC genes. Although this mutant failed to respond to the NRC4-dependent NLR Rpi-blb2, it remained unaltered in flg22-induced responses. We conclude that the NRC4 genes are not essential for flg22-induced responses in tomato.

---

Plants utilise cell surface pattern recognition receptors (PRRs) and intracellular nucleotide-binding domain leucine-rich repeat containing receptors (NLRs) to fend off invading pathogens (Dodds and Rathjen, 2010; Win et al., 2012). Both types of immune receptors detect pathogen molecules directly or indirectly to activate complex immune responses and disease resistance (Kourelis and van der Hoorn, 2018). Although PRR- and NLR-triggered immunity are generally thought to activate distinct pathways, they can induce similar outputs such as production of reactive oxygen species (ROS) and hypersensitive cell death (Peng et al., 2018). Both PRR- and NLR-activated pathways involve calcium-dependent protein kinases, mitogen-activate protein kinases (MAPK), phytohormone signalling and transcriptional reprogramming (Peng et al., 2018). However, whether these two pathways converge at some point to potentiate and strengthen the immune response remains unclear.

Throughout evolution, a subset of NLR proteins have functionally diversified into sensors that detect pathogen molecules and helpers (also known as executors) that operate genetically downstream of sensor NLRs in mediating the hypersensitive response (HR) and disease resistance (Cesari, 2018; Adachi et al., 2019). The emerging view is that although some singleton NLRs carry both activities, sensor and helper NLRs form receptor complexes that range from pairs to networks (Wu et al., 2018; Adachi et al., 2019). One example of an NLR network is formed by the NRCs (NLR-required for cell death) in asterid plants (Wu et al., 2017). Over the last ~100 million years, the NRC network has dramatically expanded from a pair of sensor and helper genes to form a complex network of phylogenetically related sensor and helper NLRs. In the *Solanaceae*, the NLR helpers *NRC2, NRC3* and *NRC4* are partially redundant but display varying degrees of specificity towards sensor NLRs that confer resistance to oomycete, bacterial and viral pathogens (Wu et al., 2017). Interestingly, a recent study linked the tomato NRC *SlNRC4a* to PRR-triggered immunity (Leibman-Markus et al., 2018b). Leibman-Markus et al. (2018) reported that overexpression of *SlNRC4a* in *Nicotiana* spp. enhances ROS production elicited by the bacterial flagellin peptide flg22 and the fungal protein ethylene-inducing xylanase (EIX). SlNRC4a associates with the PRRs AtFLS2 and LeEIX in co-immunoprecipitation experiments. Furthermore, a tomato mutant that carries a truncated 67 amino acid SlNRC4a due to a nonsense mutation introduced by CRISPR/Cas9 displays higher LeEIX-mediated ROS production and increased resistance to the necrotrophic fungal pathogen *Botrytis cinerea*. These results led Leibman-Markus et al. (2018) to conclude that SlNRC4a is a positive regulator of the immune response mediated by PRRs, notably the extensively studied FLS2 receptor.

Despite the Leibman-Markus et al. (2018) findings, the degree to which loss-of-function mutation of *NRC4* affects flg22-elicited responses has not been unambiguously determined. This experiment is somewhat limited by the observation that *SlNRC4a* (Solyc04g007070, referred to as *NRC4a* from here on) occurs in the tomato genome as a gene cluster together with two closely related paralogous genes (*SlNRC4b*, Solyc04g007060, *NRC4b*; *SlNRC4c*, Solyc04g007030, *NRC4c*), which may be functionally redundant (Supplemental Fig. S1A). This gene cluster also contains another gene Solyc04g007050, we named *SlNRC5* (*NRC5*), that is phylogenetically related to the NRCs (Wu et al., 2017). In this study, we decided to take advantage of the CRISPR/Cas9 system to generate a loss-of-function mutants in the clustered NRC genes. We reasoned that the contribution of genetically redundant *NRC4* paralogs in FLS2 mediated responses can be addressed by deleting the entire *NRC4/5* gene cluster.

To knockout the *NRC4* gene cluster of tomato, we designed four guide RNAs based on the conserved sequences in the NRC4 paralogs (Supplemental Fig. S1B). We transformed these guide RNAs together with Cas9 and kanamycin selection marker into tomato (*Solanum lycopersicum*) GCR758 (Balint-Kurti et al., 1995). We recovered 13 independent transformants that are kanamycin resistant. To determine whether these transformants are mutated in the *NRC4* gene cluster, we used gene specific primers to amplify fragments of *NRC4a*, *NRC4b*, *NRC4c* and *NRC5* (Supplemental Fig. S1C, Supplemental Table S1). These primers amplified fragments with expected sizes when genomic DNA from the wild type plants was used as template in the PCR reaction but failed to amplify some of the *NRCs* (such as *NRC4c* and *NRC4a*) with genomic DNA from the line T0-1 (Supplemental Fig. S1C). Interestingly, we could not amplify any of the *NRC4* and *NRC5* fragments from the genomic DNA of the line T0-7, suggesting that this line contained multiple deletions or a large deletion in the locus of *NRC4* gene cluster (Supplemental Fig. S1C). To further confirm the genotype of the T0-7 plant, we designed four additional primers based on the sequences adjacent to *NRC4c* and *NRC4a*. Due to the distance between the primers (over 50 kb based on the reference sequence), these primer pairs LR1F x 7075F and 7020R x 7075F could not amplify any fragments when the genomic DNA from the wild type plant was used as template (Supplemental Fig. S2). However, we successfully amplified fragments of 1.3 kb and 3.8 kb with the primer pairs LR1F x 7075F and 7020R x 7075F, respectively, using DNA from T0-7 (Fig. 1; Supplemental Fig. S2). Thus, we sequenced the 1.3 kb fragment amplified using the primer pair LR1F x 7075F by Sanger sequencing and confirmed that this plant contains a 53 kb deletion in the *NRC4* locus, connecting the open reading frame (ORF) of *NRC4c* to the ORF of Solyc04g007075 (Fig. 1; Supplemental Fig. S3). In addition to the 53 kb deletion, we also found a 290 bp deletion in *NRC4c* (Fig. 1C). The remaining sequence resulted in a fusion of ORFs from Solyc04g007075 and NRC4c with multiple frameshift mutations leading to premature stop codons in *NRC4c* (Supplemental Fig. S3). We further obtained a homozygous T2 line (*nrc4*_7.4) and used this line for further experiments (Supplemental Fig. S4).

**Figure 1.**
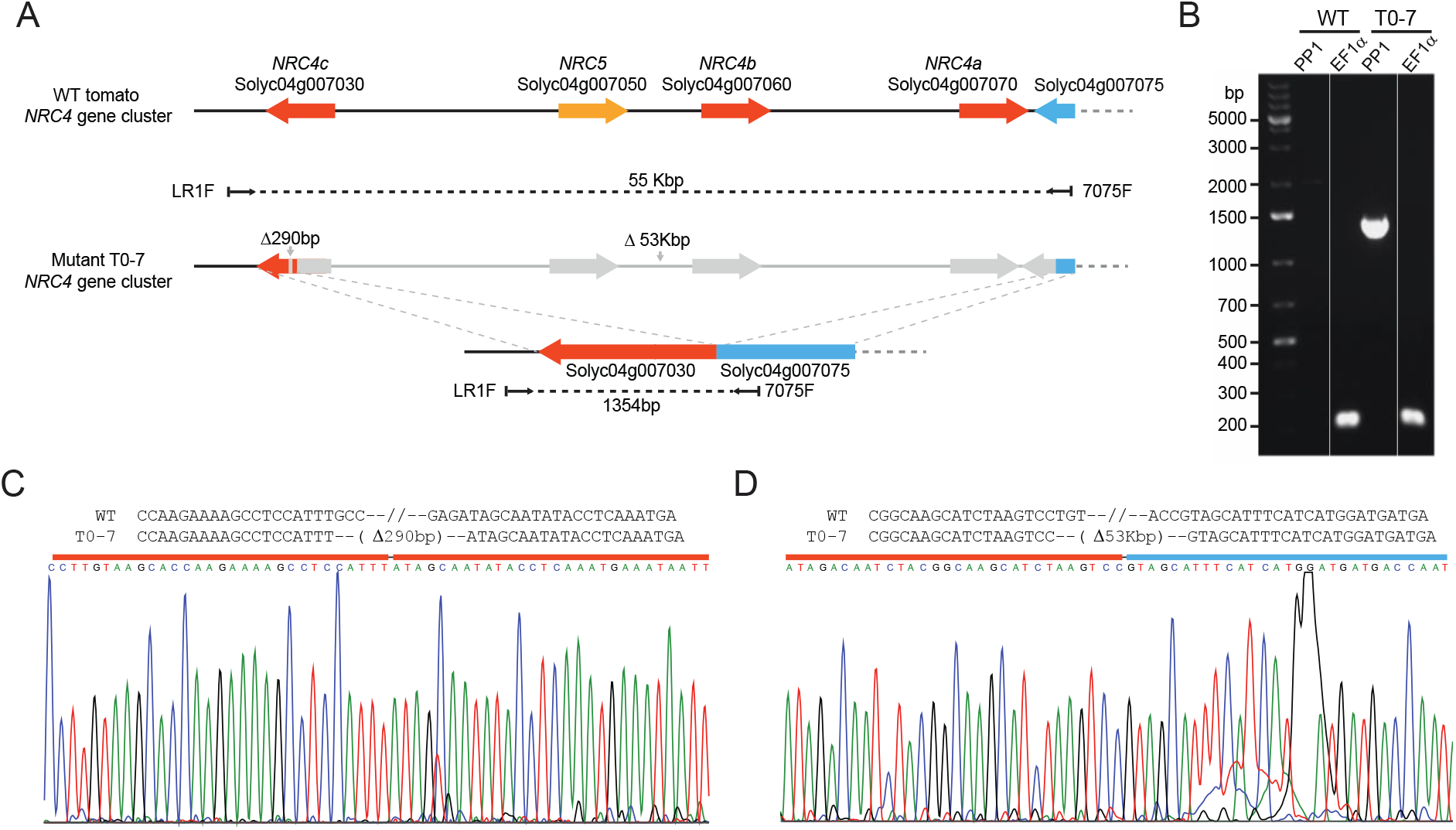
The T0-7 transformant carries a large (>53 kb) deletion spanning across the NRC4 gene cluster. A. Schematic view of the tomato NRC4 gene cluster in wild-type (WT) and mutant T0-7. Orange, NRC4 paralogs; yellow, NRC5; blue, Solyc04g007075, which contains incomplete sequence information due to a sequencing gap in the reference genome. The deleted regions in the mutant T0-7 are marked in grey. B. PCR-genotyping for the large deletion. PP1, amplification with primer pair 1:LR1F x 7075F indicated in A; EF1α, amplification control with EF1α primers. The uncropped image is provided in Supplemental Fig. S2B. C. Sequence alignment and chromatograms of Sanger DNA sequencing results. In this region, the mutant T0-7 contains a 290bp deletion based on reference genome and the results of sequencing. D. Sequence alignment and chromatograms of Sanger DNA sequencing results. In this region, the mutant T0-7 contains a 53kbp deletion based on the reference genome and the results of sequencing.

We previously reported that the sensor NLR Rpi-blb2, which confers resistance to potato late blight pathogen *Phytophthora infestans*, depends on NRC4 when expressed in *N. benthamiana*. To test whether Rpi-blb2 signals through NRC4 in tomato, we expressed Rpi-blb2/AVRblb2, Rpi-vnt1/AVRvnt1 (NRC-independent) or GFP in wild type and the *NRC4* knockout tomato line using agroinfiltration (see Supplemental methods). Rpi-blb2-mediated cell death was compromised in the *NRC4* knockout plants, whereas Rpi-vnt1 triggered strong cell death in both wild type and *NRC4* knockout plants, consistent with the earlier finding from the *N. benthamiana* experimental system (Supplemental Fig. S4).

Leibman-Markus et al. (2018) proposed that *NRC4a* participates in immunity mediated by FLS2 because overexpression of *NRC4a* in *N. benthamiana* enhances ROS production after flg22 treatment. Leibman-Markus et al. (2018) obtained a CRISPR/Cas9 mutagenized tomato line that expresses a truncated variant of SlNRC4a. However, the effect of this mutation on FLS2 responses was not reported. As *NRC4a* exists in a gene cluster with the highly homologous *NRC4b* and *NRC4c* that are potentially functionally redundant, we reasoned that our *NRC4/5* gene cluster deletion would be ideal to test whether the NRC4 genes are required for FLS2-mediated responses. To test the hypothesis, we monitored apoplastic ROS production in response to flg22 peptides. We observed a transient flg22-induced ROS burst with the leaf discs from wild type tomato plant GCR758. However, we did not observe a notable difference in terms of flg22-induced ROS burst between the wild type and the *NRC4* deletion line *nrc4*_7.4 (Fig. 2A and B). As mitogen-activated protein kinase (MAPK) activation represents another typical output in FLS2-mediated responses, we tested whether the MAPK phosphorylation was impaired in the *NRC4* knockout plants. We detected increased phosphorylation of MAPKs by Western blotting with p-42/44 antibody after flg22 treatment. We detected increased phosphorylation of MAPKs in the *NRC4* knockout mutant and, here too, we did not observe a significant difference between the wild type and *NRC4* deletion mutant. Our results indicate that the *NRC4* genes are not essential for flg22-induced responses in tomato (Fig. 2C, Supplemental Fig. S5).

**Figure 2.**
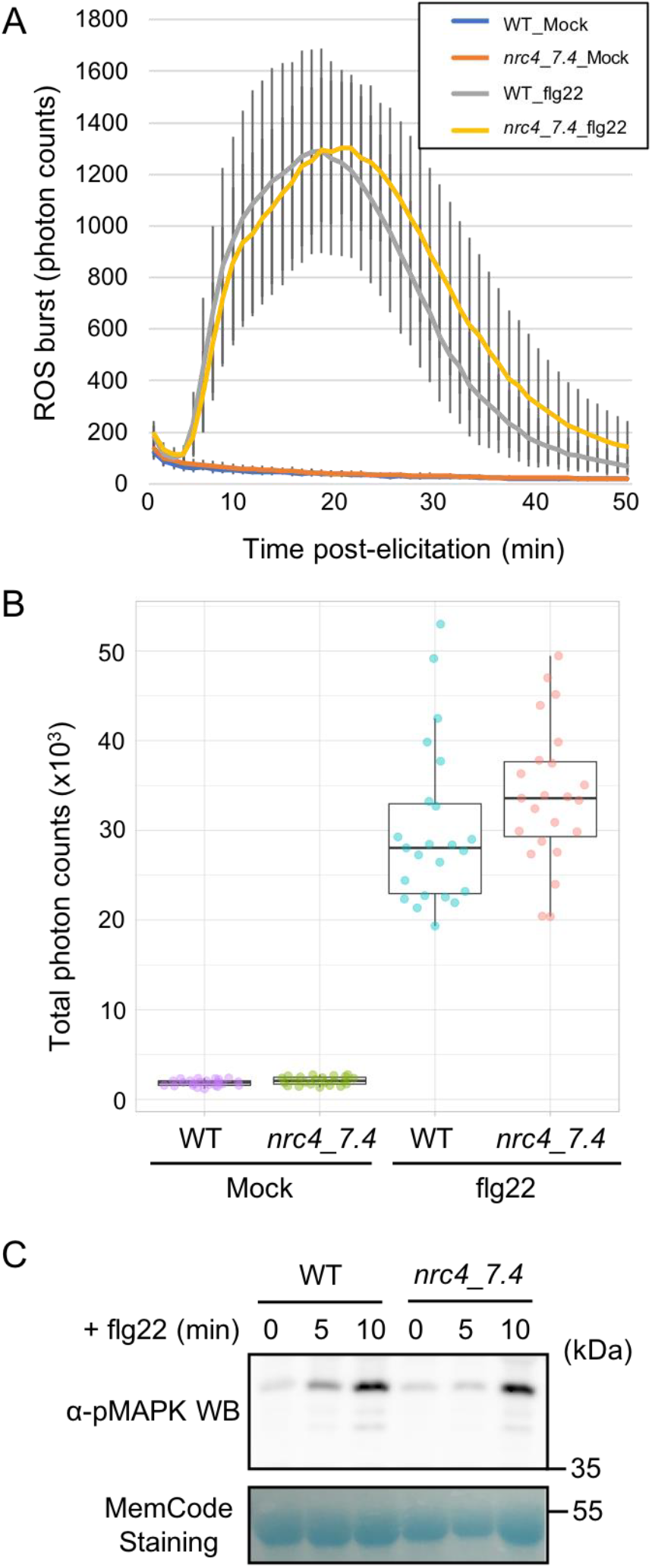
NRC4 knockout tomato plants are not impaired in FLS2-mediated defence responses. A and B. Flg22-triggered ROS bursts were measured for 50 min using leaf discs of the WT and T2 line *nrc4*_7.4 (A). Total photon counts of each treatment during 50 min were used for scatter plot analyses (B). C. Flg22-triggered MAPK activation was analysed by immunoblots with α-pMAPK. Proteins were extracted from tomato leaf tissues of the WT and T2 line *nrc4*_7.4, 0, 5 or 10 min after treatment with flg22.

Previous studies have suggested that the NRC proteins are involved in immune responses mediated by both intracellular NLR and cell surface PRR immune receptors (Gabriels et al., 2007; Wu et al., 2016; Brendolise et al., 2017; Wu et al., 2017). Silencing of *NRC2* and *NRC3* by virus-induced gene silencing (VIGS) and RNAi reduces Cf4- and Prf-mediated cell death in *N. benthamiana*, indicating that NRC2 and NRC3 are involved in cell death responses activated in both pathways (Brendolise et al., 2017; Wu et al., 2016). However, the results we obtained here with our *NRC4* knockout line raise questions about the precise role of *NRC4a* in FLS2-mediated responses (Leibman-Markus et al., 2018a; Leibman-Markus et al., 2018b). Although we cannot rule out that other NRC homologs, such as NRC2 and NRC3, contribute to FLS2-mediated ROS production, we failed to observe any measurable differences between the wild type and the *NRC4* deletion line in our flagellin immunoassays.

The NRC network is phylogenetically restricted to asterids and caryophyllales but missing in Arabidopsis and other rosid species (Wu et al., 2017). Therefore, our results may not be that surprising given that FLS2 belongs to an ancient receptor-like kinase subfamily XII that broadly occurs in angiosperms (Dufayard et al., 2017; Liu et al., 2017). In contrast, NRCs may be involved in Cf-4 and LeEIX mediated immunoresponses considering that these cell surface receptor-like proteins are phylogenetically restricted to some asterid clades (Kang and Yeom, 2018) and, unlike flg22, trigger hypersensitive cell death in plant tissues (Gabriels et al., 2007; Brendolise et al., 2017; Wu et al., 2016). Future work will need to address how cell surface receptors mechanistically engage NLR proteins to induce cell death and other immune responses.

## Supplemental Data

The following supplemental materials are available.

Supplemental Methods

**Supplemental Figure S1.**
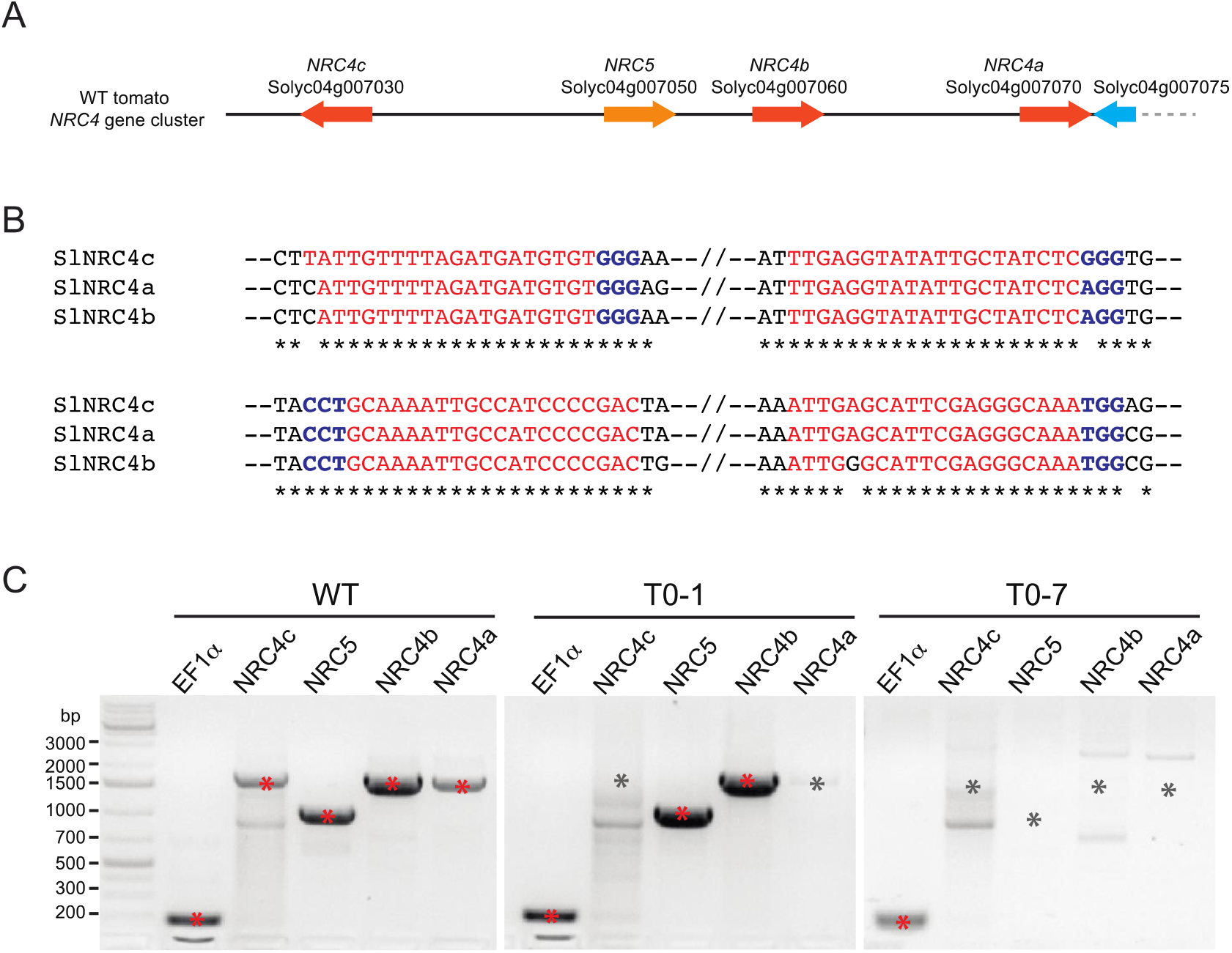
Targeting the NRC4 gene cluster with CRISPR/Cas9 in tomato. A. Cartoon showing the NRC4 gene cluster (Solyc04g007030, Solyc04g007060 and Solyc04g007070) targeted by CRISPR/Cas9 in this study. Orange, NRC4 paralogs; yellow, NRC5; blue, Solyc04g007075, which contains incomplete sequence information due to a sequencing gap in the reference genome. B. Sequence alignment of the region targeted by sgRNA in the NRC4 paralogs. The PAM motifs are marked in blue, and the sgRNA sequences are marked in red. C. PCR-genotyping for the presence of NRC4 and NRC5 genes in T0 transgenic lines. Amplification bands with expected size are labelled with red asterisks, whereas missing bands at the corresponding size are labelled with grey asterisks.

**Supplemental Figure S2.**
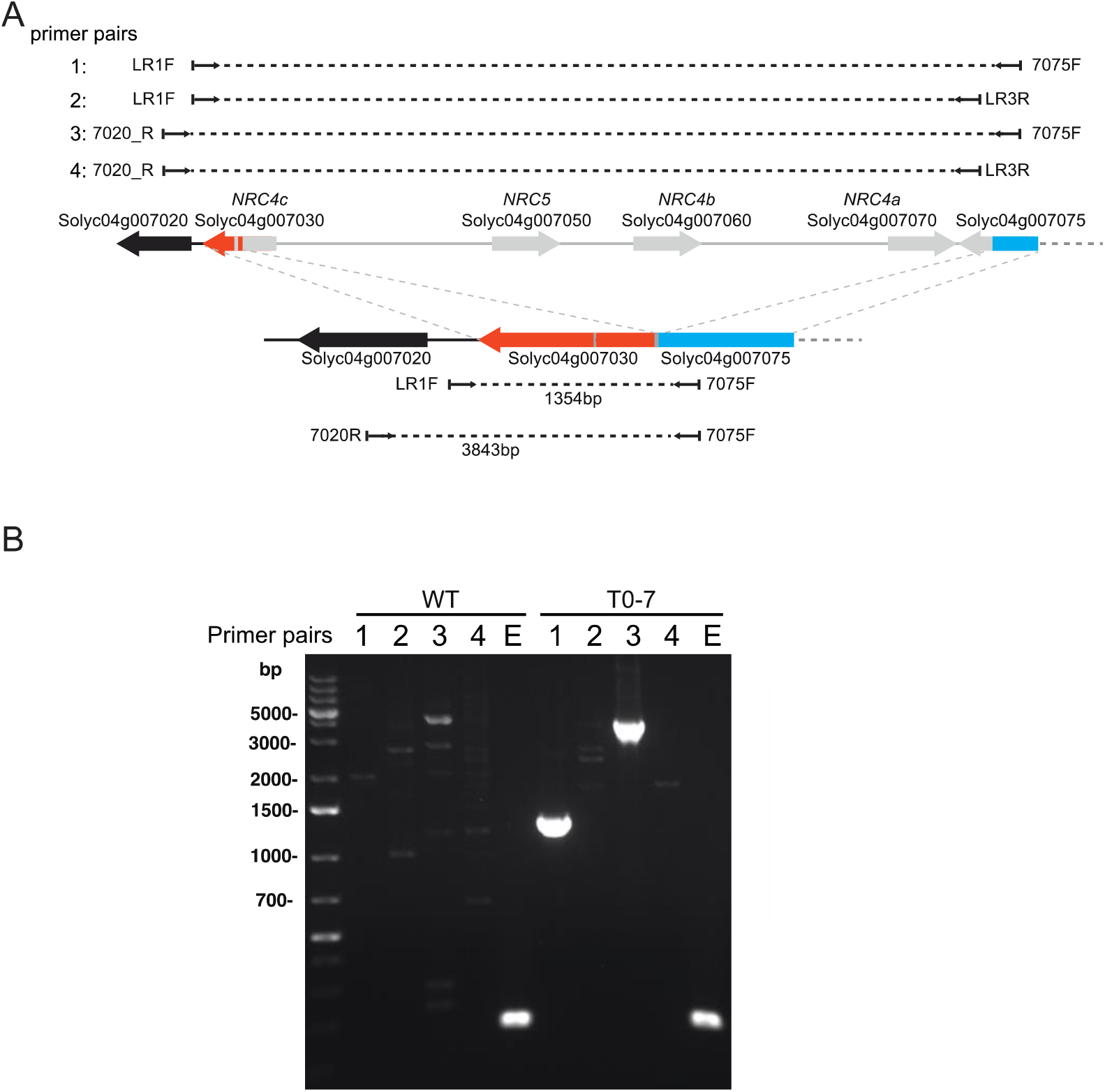
Primer design and characterisation of the large deletion in mutant T0-7. A. Cartoon showing the NRC4 gene cluster and positions of primers for characterising the mutant T0-7. B. Agarose gel electrophoresis of the PCR products obtained with primer pairs indicated in A. None of the primer pairs amplified fragments with expected size when the genomic DNA from the WT plant was used. Primer pairs 1:LR1F x 7075F and 3:7020R x 7075F amplified fragments of 1354bp and 3848bp with the genomic DNA from mutant T0-7. Primer pairs 2: LR1F x LR3R and 4: 7020R x LR3R failed to amplify any fragments because of the reverse primer LR3R locates on the region that is deleted in the mutant T0-7. A primer pair for EF1α (E) was used as amplification control.

**Supplemental Figure S3.**
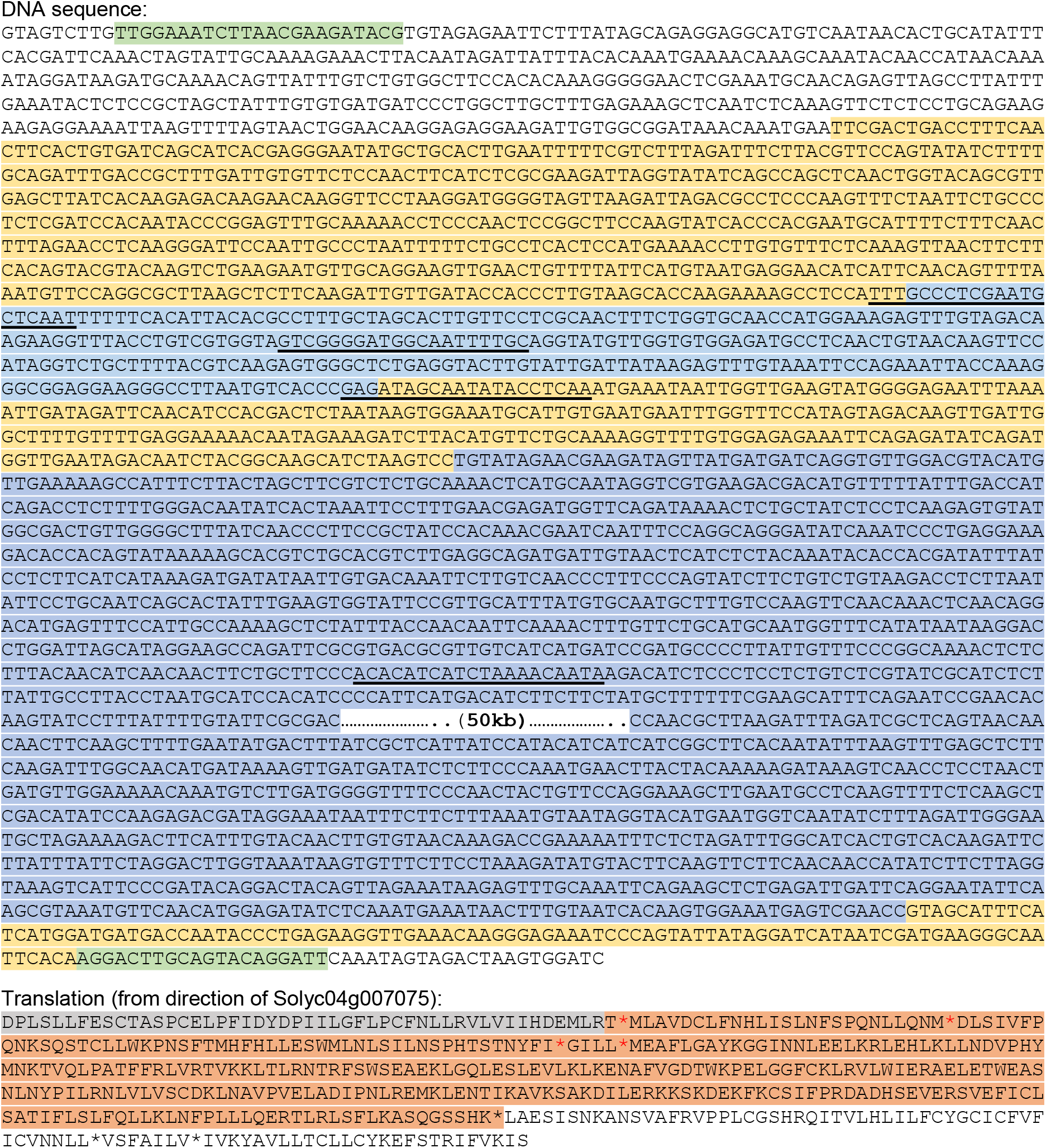
Sanger sequencing result of the NRC4 deletion allele from T0-7. In the DNA sequence, positions of the primers LR1F and 7075F used to amplify across the deleted region are highlighted in green. Regions of the NRC4 gene cluster present in the PCR amplicon are highlighted in yellow. Positions of the sgRNAs targets are underlined. Deleted regions are shown in blue. In the translation, sequence belonging to Solyc04g007075 is marked in grey, and sequence belonging to NRC4c (Solyc04g007030) is marked in orange. Red asterisks represent stop codons.

**Supplemental Figure S4.**
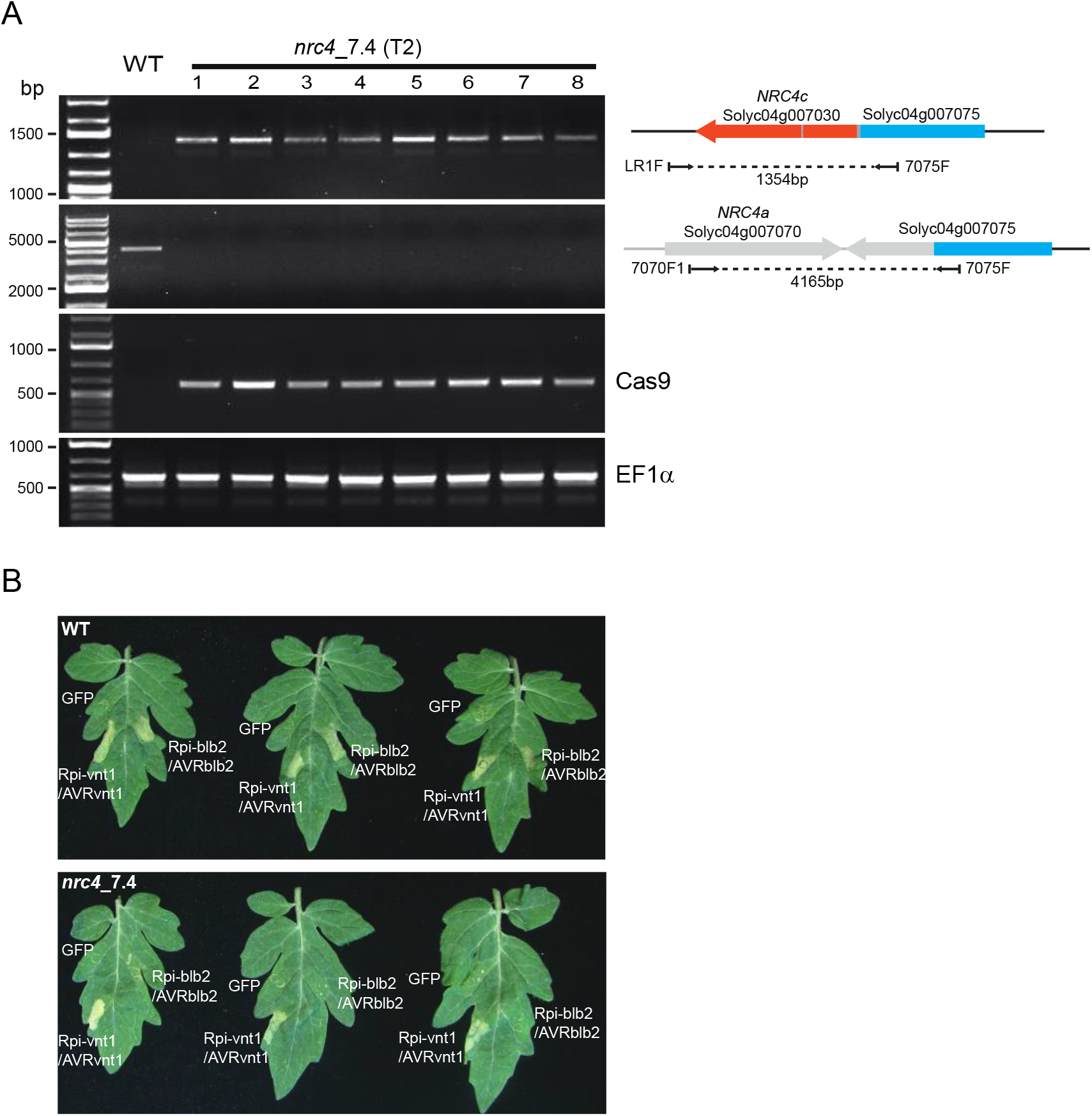
The NRC4 knockout homozygous T2 line (*nrc4*_7.4) is impaired in Rpi-blb2 mediated cell death. A. Genotyping of the T2 line *nrc4*_7.4. Primer pair LR1F x 7075F and primer pair 7070F1 x 7075F were used to characterise the NRC4 gene cluster in the WT and *nrc4*_7.4 plants. Eight T2 plants from the line *nrc4*_7.4 were used. Primers for amplification of Cas9 and EF1α were used as controls. B. Cell death assay on leaves from WT and *nrc4*_7.4 plants. Transient expression of Rpi-blb2 and AVRblb2 induced cell death in WT, but not in *nrc4*_7.4 leaves. Constructs expressing GFP, Rpi-vnt1 and AVRvnt1 were used as controls.

**Supplemental Table S1.**
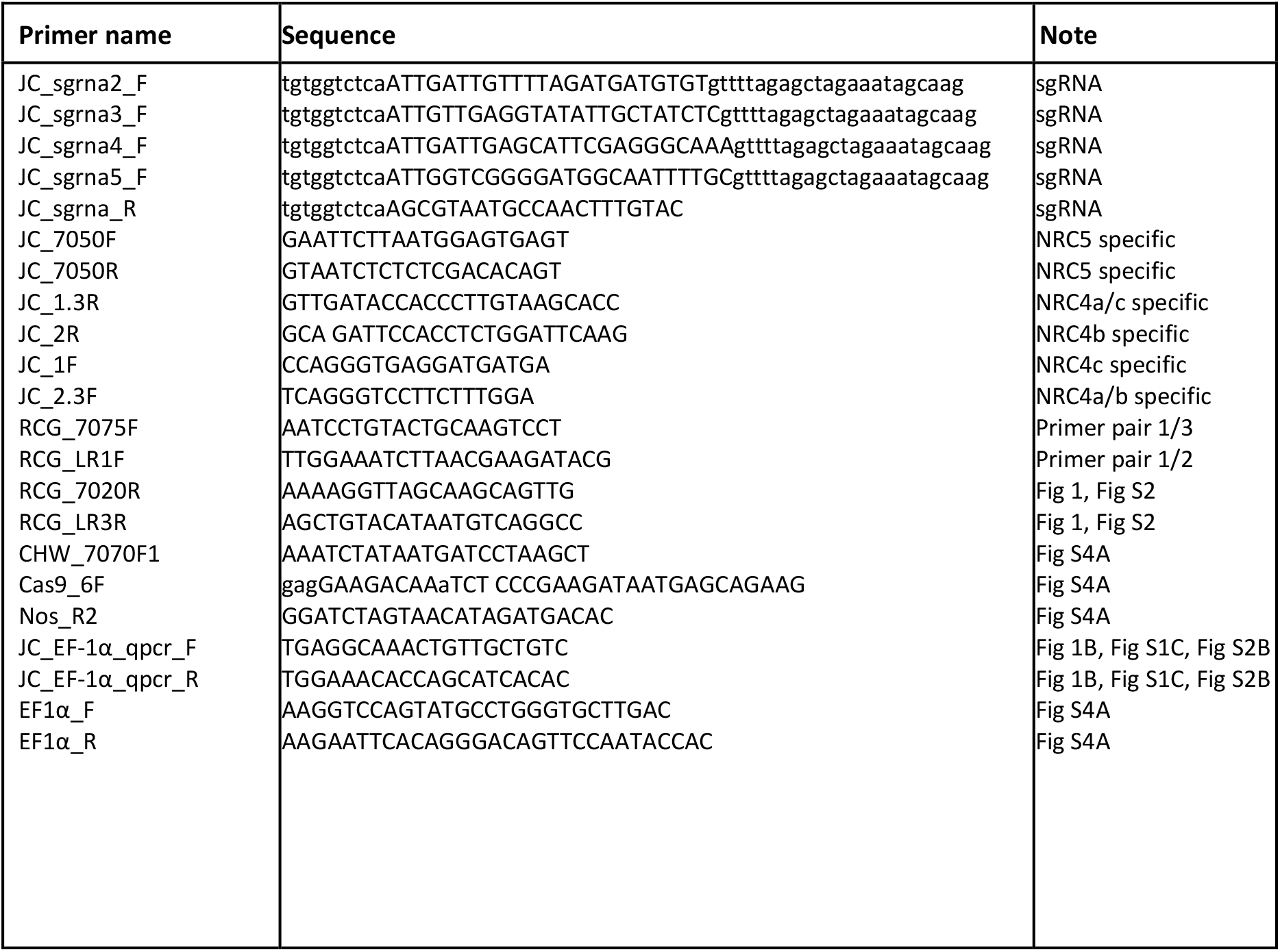
Primers used in this study.

## ACKNOWLEDGEMENTS

We thank the Tissue Culture and Transformation Team at The Sainsbury Laboratory for performing tomato transformation, Marta Bjornson for helping with the ROS assays, Bruno Ngou and Hailong Guo for helping with the MAPK phosphorylation assays.

## Supplemental Data

### Supplemental Methods

#### CRISPR/Cas9 Construct Assembly

All constructs were assembled using the Golden Gate cloning method (Weber et al. 2011). Level 1 constructs carrying sgRNAs placed under the control of the Arabidopsis (*Arabidopsis thaliana*) U6 promoter were assembled as described (Belhaj et al. 2013). Primers JC_sgrna2_F, JC_sgrna3_F, JC_sgrna4_F and JC_sgrna5_F forward primers were used in combination with the JC_sgrna_R reverse primer (Supplemental Table S1) to clone sgRNAs 2, 3, 4 and 5, respectively.

Level 1 constructs pICH47732::NOSp::NPTII (addgene no. 51144), pICH47742::35S::Cas9 (addgene no. 49771), pICH47751::AtU6p::sgRNA2, pICH47761::AtU6p::sgRNA3, pICH47772::AtU6p::sgRNA4, pICH47781::AtU6p::sgRNA5 and the linker pICH41822 (addgene no. 48021) were assembled into the level 2 vector pAGM4723 (addgene no. 48015) as described (Weber et al. 2011) resulting in the level 2 construct pAGM4723::NPTII::Cas9::sgRNA2::sgRNA3::sgRNA4::sgRNA5.

#### Plant transformation

Tomato cultivar GCR758 (Balint-Kurti et al. 1995) was transformed with the pAGM4723::NPTII::Cas9::sgRNA2::sgRNA3::sgRNA4::sgRNA5 construct as previously described (Fillatti et al. 1987). T0 transgenic plants were selected in the medium with kanamycin and then transferred into soil.

#### Plant genotyping

Genomic DNA was extracted from T0 transformants as described (Nekrasov et al. 2017). Plants were genotyped using PCR with respective primers (Supplemental Table S1).

#### Measurement of ROS

ROS production after flg22 peptide treatment was monitored by a luminol-based assay. 4 mm diameter tomato leaf discs were incubated for 16 hours in 150 µL of water in a 96-well plate. Before measurement of ROS, the water was removed and 100 µL of ROS detection solution [17 mM luminol (Sigma); 1 µM horseradish peroxidase (Sigma); 100 nM flg22 (EzBiolab)] was added to each well. Chemiluminescence was measured by using an ICCD photon-counting camera (Photek, East Sussex, UK).

#### Detection of MAPK phosphorylation

Six leaf discs (8 mm diameter) of tomato leaves and were homogenised in 200 µL of extraction buffer [10% glycerol; 25 mM Tris, pH 7.5; 1 mM EDTA; 150 mM NaCl; 2% w/v PVPP; 10 mM DTT; 1% v/v protease inhibitor cocktail 2 (Sigma, P5726); 1% v/v protease inhibitor cocktail 3 (Sigma, P0044); 0.2% v/v IGEPAL (Sigma). After centrifugation at 12,000 *xg* for 10 min, 50 µL of supernatant was mixed with equal amount of 2x SDS-PAGE sample buffer and used for SDS-PAGE. Immunoblotting was done with an antibody against phosphorylated MAPK (Phospho-p44/42 MAPK Erk 1/2 Thr202/Tyr204 XP Rabbit mAb #4370; Cell Signaling Technology).

#### Cell death assay on tomato leaves

Transient expression of GFP, Rpi-blb2, AVRblb2, and Rpi-vnt1 and AVRvnt1 in tomato leaves were performed according to the method described previously by Bos et al. (2006). Suspensions of *Agrobacterium tumefaciens* AGL1 strains harboring the expression vector of different proteins were prepared in infiltration buffer (10mM MES, 10mM MgCl2, and 150μM acetosyringone, pH5.6) and adjusted to the final OD_600_ of 0.3 for each expression construct. Leaflets of the 4th leaves from four-week-old tomato plants were syringe-infiltrated with *A. tumefaciens* strains harboring the expression vector of different proteins as indicated.

